# Uneven distribution of Purkinje cell injury in the cerebellar vermis of term neonates with hypoxic-ischemic encephalopathy

**DOI:** 10.1101/2020.04.15.043018

**Authors:** K.V. Annink, I.C.E. van Leeuwen, N.E. van der Aa, T. Alderliesten, F. Groenendaal, R.K. Jellema, C.H.A. Nijboer, P.G.J. Nikkels, M. Lammens, M.J.N.L. Benders, F.E. Hoebeek, J. Dudink

**Author notes:** Corresponding author: J. Dudink. Full corresponding address: Wilhelmina Children’s Hospital, room number KE04.123.1, Lundlaan 6, 3508 AB Utrecht, P.O. Box 85090, the Netherlands. shared first author (K.V. Annink, I.C.E. van Leeuwen). shared last author (F.E. Hoebeek, J. Dudink).

## Abstract

**Introduction:** In term neonates with hypoxic-ischemic encephalopathy (HIE), cerebellar injury is becoming more and more acknowledged. Animal studies demonstrated that Purkinje cells (PCs) are especially vulnerable for hypoxic-ischemic injury. In neonates, however, the extent and pattern of PC injury has not been investigated. The aim of this study was to determine the distribution of PC injury in the cerebellar vermis of term born neonates with HIE.

**Methods:** Term born neonates with HIE that underwent post-mortem autopsy of the cerebellar vermis were included. Haematoxylin & Eosin (H&E) stained sections of the vermis were used to determine total PC count and morphology (normal, abnormal or non-classified) at the bases and crown of the folia and of the lobules in both the anterior and posterior lobes. Differences in PC count and PC morphology between the anterior and posterior lobe and between the bases and crown were calculated using the paired samples T-test or Wilcoxon-signed rank test.

**Results:** The total number of PCs were significantly higher at the crown compared to the bases (p<0.001) irrespective of the precise location. Besides, PCs at the bases more often had an abnormal morphology. No significant difference between the total number of PCs in the anterior and posterior lobe was observed.

**Conclusion:** The abnormal PC count and morphology in term neonates with HIE resembles supratentorial ulegyria.

## INTRODUCTION

Hypoxic-ischemic encephalopathy (HIE) following perinatal asphyxia is the leading cause of mortality in term neonates with an incidence of 1.5 per 1000 term births (1,2). Therapeutic hypothermia is the current standard of care to reduce brain injury and additional treatment options are still limited (3). Besides high mortality rates, long-term neurodevelopmental disabilities such as cognitive, behavioural and attention impairments and motor dysfunction are still common in neonates with moderate or severe HIE (4–7).

In the clinical setting, HIE is often characterized as cerebral injury (8). However, also cerebellar injury in neonates is being increasingly recognized as a contributor to neurodevelopmental outcome (9,10). For instance, cerebellar injury has been linked to motor deficits, as well as cognitive functioning, behaviour, learning and emotional deficits (11,12). However, research about the association between behavioural problems and cerebellar injury in HIE remains sparse (9). In infants born extremely preterm (i.e. below 28 weeks of gestation) a clear association was found between early cerebellar injury and long-term behavioural problems (10,13).

During the third trimester of pregnancy the cerebellum shows a rapid increase in growth, making the cerebellum especially vulnerable to insults that disturb normal development within this time period (e.g. hypoxia-ischemia) (14,15). Surprisingly, acute cerebellar injury is rarely detected on conventional clinical neuroimaging (e.g. T1-, T2-weighted and diffusion weighted MRI scans) (16,17). In contrast, on histopathological level, there is evidence that the cerebellum is injured after severe HIE in infants (18). Of the cerebellar neurons, the Purkinje cells (PCs) are known to be the most susceptible to hypoxic events, which is most likely due to their high metabolic activity and associated oxygen demand (19,20). Post-mortem studies in human neonates with HIE on PCs are lacking, but in neonatal mouse models, pronounced reductions in PC number and morphological alterations of PCs in response to hypoxic-ischemia were indeed found (i.e. swelling, autolytic necrosis, cell shrinkage and dark cell degeneration) (24,25). Other animal studies on perinatal hypoxic-ischemia have also demonstrated increased microglia activation, myelination deficits and overall necrosis and apoptosis in the cerebellum (23–25).

Whereas the cerebellar anatomy has been described as homogenous, its susceptibility to cellular damage has been shown to vary between cerebellar lobules. In murine models, it has been shown that PCs in lobules IX and X were less vulnerable for hypoxic-ischemia (26). Additionally, the lobes of the adult human cerebellum have a functional topographic organization, so the pattern of PC injury might be relevant for functional outcome (25, 26). Literature about the topographic organization of the vermis in neonates with HIE is scarce as well as literature about differences in vulnerability of the vermis to hypoxia. Therefore, the aim of this study was to determine the distribution of PC injury in the vermis of term born neonates with HIE.

## PATIENTS AND METHODS

### Patient population

In this retrospective cohort study, (near)-term neonates (36 until 42 weeks of gestational age) with HIE were included, who were treated with therapeutic hypothermia (born between 2008 and 2015) or would have qualified for therapeutic hypothermia according to the criteria of Azzopardi *et al*. (born between 2000 and 2008) and who underwent autopsy after neonatal death with haematoxylin and eosin (H&E) staining of the vermis (3). Exclusion criteria were perinatal asphyxia based on major congenital abnormalities, metabolic disorders, chromosomal abnormalities or poor quality of the sections of the vermis. No normotypic age-matched controls were available in this study.

Parental consent was obtained for post-mortem histopathological examination. The Medical Ethical Committee (MEC) of UMC Utrecht confirmed that the ‘Medical Research Involving Human Subjects Act’ did not apply to this study and therefore an official approval of the MEC was not required (MEC # 18-167). The UMC Utrecht biobank approved the use of the rest tissue (biobank # 18-284).

### Histopathological examination

Brains were removed during autopsy and before the slices could be cut they were dehydrated with ethanol 70%. Afterwards they were fixed in a 4% buffered formalin solution for 4-6 weeks (18). The vermis was cut midsagittally in 5-6 µm thick sections, mounted on coated slides and stained for H&E. From each patient one section was implemented.

The H&E stained sections were used to examine the presence and the morphology of PCs using the light microscope Zeiss AXIO lab A1 and ZEN 1.2 lite software (Zeiss, CA, U.S.A). First, the sections of the vermis were photographed at various locations with 63x magnification (Figure 1). For each patient the pattern of PC injury was examined at two levels: (1) within the lobules (locations examined: the bases of the sulci and crown of lobules II, III, VIII, IX), and (2) within two folia (locations examined: the bases of the sulci and crown of a proximal folia and a distal folia of lobules II, III, V, VI, VIII, IX). The crown is defined as the most convex part of a folia or lobe (Figure 1B and 2A) and the base as the most concave part. The base of a sulcus is the mean value of both sides of the base. The anterior values are the average values of lobules II, III and V and the posterior values are the average of lobules VI, VIII and IX. However, data were only included throughout examination when all values within one lobule were available. Whenever lobular data were available, but data from the folia were missing (or vice versa), only lobular data were included since this was separately investigated.

**Figure 1:**
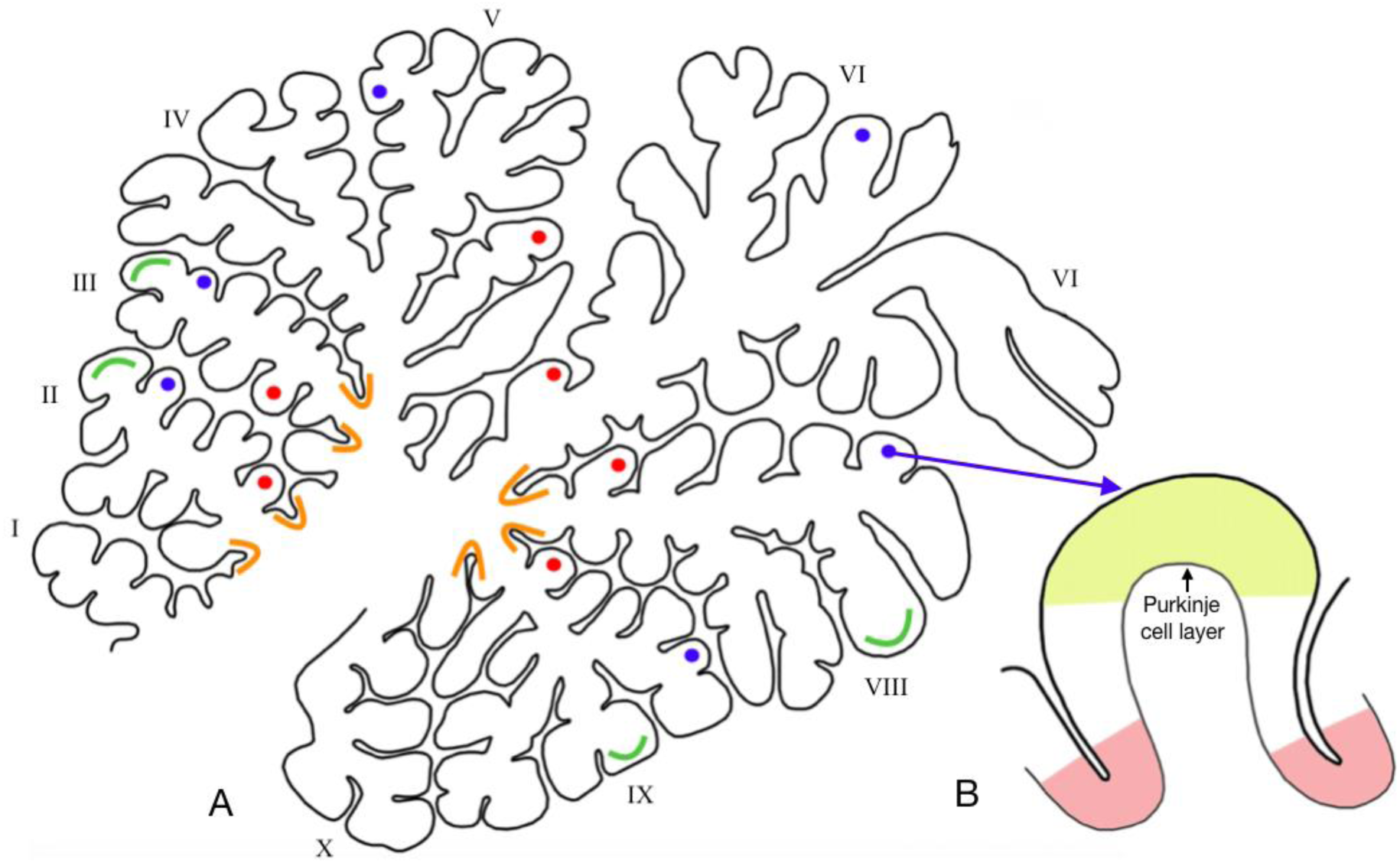
Locations photographed for analyses in the vermis. A. The red dots represent the proximal located folia in the anterior and posterior lobes. The blue dots represent the distal located folia in the anterior and posterior lobes. The orange (concave) curvatures represent the bases of the sulci of the anterior and posterior lobules, the green (convex) curvatures represent the crowns of the anterior and posterior lobules. B. Magnified folia. The yellow bow represents the crown, the pink curvatures represent the bases of the sulci of the folia.

### PC classification system

PCs were counted and categorized at the above mentioned locations in the vermis using a newly developed PC classification system based on Hausmann *et al*. (19) by an experienced neuropathologist (M.L.). PCs were only counted when located in the Purkinje cell layer and when a clear nucleus was visible. PCs were categorised as: normal, abnormal or non-classified (Table 1 and Figure 2). The numbers of normal, abnormal and non-classified PCs were counted for each location using ImageJ software version 1.47 (Wayne Rasband, NIH, USA). Afterwards the numbers were corrected for the measured distance in cells per 100 µm. Also the percentage of abnormal PCs compared to normal PCs was calculated for every location.

**Table 1:** Purkinje cell classification system.

| Category | Definition |
| --- | --- |
| Normal | 1. An intact cell membrane<br>2. A basophilic/light stained cytoplasm<br>3. A clearly identifiable, basophilic/light stained nucleus with intact nuclear membrane<br>4. An identifiable nucleolus in the nucleus<br>*(In addition to 1 and 2, criteria 3 or 4 should be met to score a PC as normal)* |
| Abnormal | 1. No intact/ruptured cell membrane<br>2. A hypereosinophilic stained cytoplasm<br>3. An identifiable nucleolus but no clear nucleus or rest of nucleolus<br>*Presence of activated Bergmann glia and shrinkage of total cell volume was taken into account as an additional confirmation of abnormality, but was not required* |
| Non-classified | 1. The nucleolus and nucleus were not visible or could not be categorized<br>2. The soma of the PC was clearly visible<br>3. The cell is located in the PC layer |

**Figure 2:**
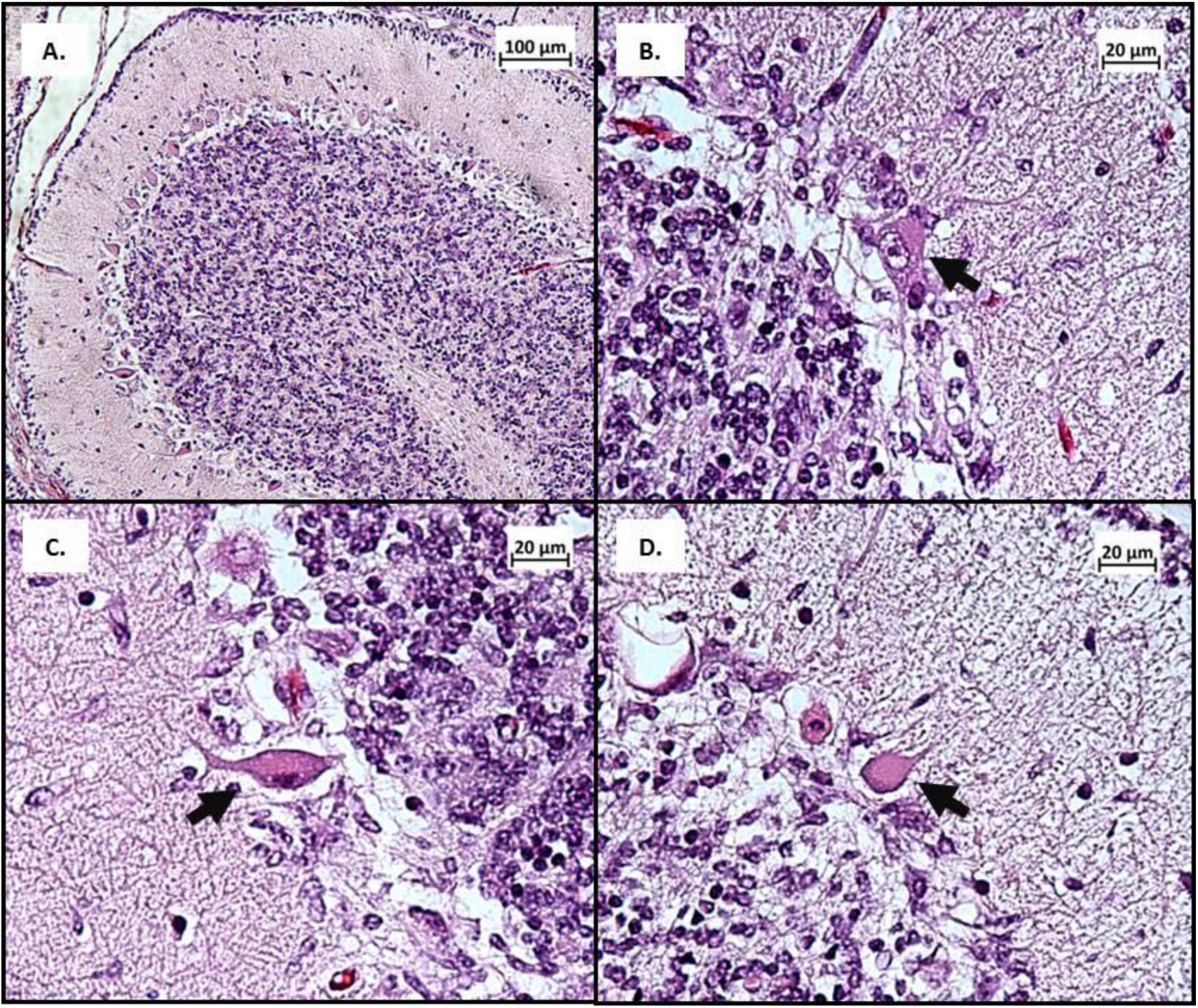
Examples of PC classification system. A. Crown of a proximal located folia of lobe 2. B. Normal PC. C. Abnormal PC. D. Non-classified PC because no nucleus or nucleolus is visible.

#### Statistical analyses

Statistical analyses were performed using IBM SPSS Statistics for Windows, version 25 (IBM New York 2013). Differences in PCs in the anterior and posterior lobe were calculated using the paired samples *t*-test for normally distributed data and a Wilcoxon signed rank test for data that were not normally distributed. Differences in the number and percentage of PCs at the crown compared to the bases of the sulci within the lobules and within the proximal and distal folia were assessed using a paired samples *t*-test for normally distributed data and a Wilcoxon signed rank test for data that were not normally distributed. *P <* 0.05 was considered significant. All data were corrected for multiple comparison analyses by multiplying the P-value with the number of comparisons per statistical test.

## RESULTS

Between 2000 and 2015, 22 patients with HIE underwent autopsy and had good quality sections of the vermis stained for H&E available for examination. Clinical data are presented in table 2. All patients died from irreversible brain injury after redirection of care. Three infants were treated with hypothermia for less than 72 hours, because therapeutic hypothermia was stopped based on clinical decision making. Of the included patients, 9 had missing values for some of the lobules. This is also specified in Table 1.

**Table 2:** patient characteristics.

| **Patient characteristics** |  |
| --- | --- |
| Gender (male), n (%) | 15 (68.2%) |
| Gestational age (weeks), median (IQR) | 40.0 (1.45) |
| Mode of delivery, n (%) |  |
| *Emergency caesarean section or fetal indication* | 11 (50.0%) |
| *Normal vaginal delivery* | 9 (40.9%) |
| *Vacuum extraction* | 2 (9.1%) |
| Apgar score at 1 min, median (IQR) | 1 (0 – 2.5) |
| Apgar score at 5 min, median (IQR) | 3 (0 – 4.5) |
| Umbilical cord pH, median (IQR) | 6.9 (0.1) |
| Birth weight (grams), median (IQR) | 3352 (429) |
| Hypothermia, n (%) |  |
| *No* | 5 (22.7%) |
| *Yes, incomplete (>48h)* | 3 (13.6%) |
| *Yes, complete (72h)* | 14 (63.6%) |
| Postnatal age at death (days), median (IQR) | 5 (3 – 5.25) |
| Time of autopsy after death (days, median (IQR) | 1 (1-2) |
| Missing values, n (%) |  |
| Anterior lobes |  |
| *Lobe II* | 9 (40.9%) |
| *Lobe III* | 9 (40.9%) |
| *Lobe V* | 4 (18.2%) |
| Posterior lobes |  |
| *Lobe VI* | 2 (9.1%) |
| *Lobe VIII* | 2 (9.1%) |
| *Lobe IX* | 3 (13.6%) |

No significant differences were detected between the total number of PCs, i.e. normal, abnormal and non-classified PCs combined, in the anterior and posterior lobes (Table 3). For all analysed locations, the total number of PCs was significantly higher at the crowns compared to the bases of the sulci (Figure 3). The percentage abnormal PCs was significantly higher at the bases of the sulci than in the crowns in the posterior lobules (p=0.011), as within the central folia (p=0.001) and peripheral folia (p=0.002) of the posterior lobules. In the anterior locations, the increase in abnormal PCs was only significant within the central folia of the anterior lobules (p=0.008).

**Table 3:** Total PC count per 100 µm in anterior versus posterior lobes.

| **Difference in PC number anterior and posterior** |  |  |
| --- | --- | --- |
| **Location** | **Total PC count anterior vs. posterior** | **P value** |
| Anterior vs. posterior lobules |  |  |
| All crowns | 0.83 (0.72 – 1.01) vs. 1.29 (1.07 – 1.51) <sup>#</sup> | 0.08 |
| All bases of sulci | 0.44 (0.27) vs. 0.48 (0.26) * | 0.33 |
| Central folia of anterior vs. posterior lobules |  |  |
| All crowns | 1.08 (0.24) vs. 0.86 (0.31) * | 0.14 |
| All bases of sulci | 0.46 (0.18) vs. 0.43 (0.22) * | $\approx 1.00$ |
| Peripheral folia of anterior vs. posterior lobules |  |  |
| All crowns | 0.99 (0.27) vs. 1.11 (0.22) * | 0.68 |
| All bases of sulci | 0.56 (0.17) vs. 0.56 (0.20) * | $\approx 1.00$ |

*The difference in total PC count in 100 $\mu\text{m}$ between lobules II, III, V (anterior) and lobules VI, VIII, IX*
*(posterior) of the vermis are presented. The paired samples t-test for normally distributed data and a Wilcoxon signed rank test for non-Gaussian data was used. The crowns and bases of the sulci of the proximal and distal folia anterior/posterior are taken together. <sup>#</sup>Median (IQR), \*Mean (SD).*

**Figure 3:**
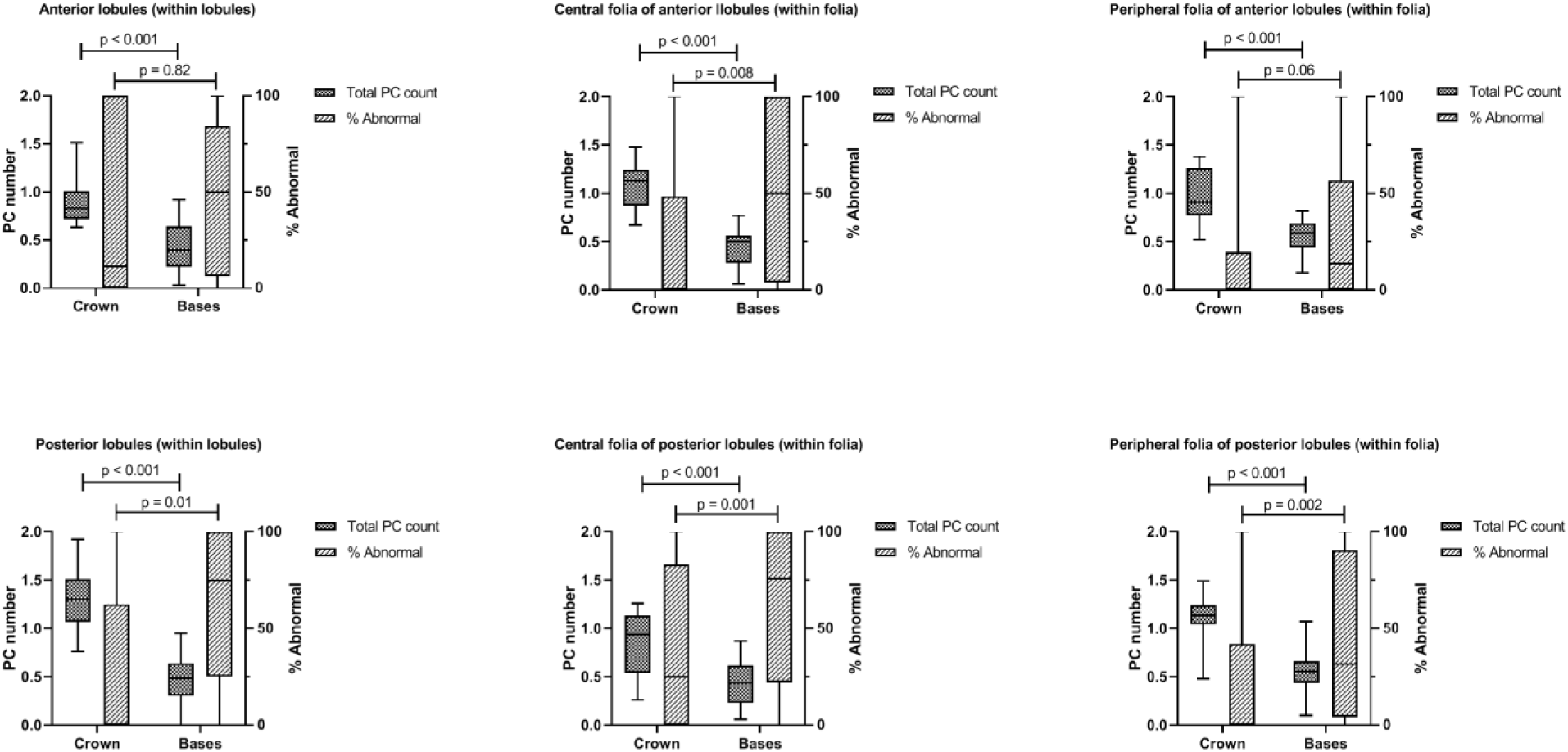
differences in PC number and abnormalities within lobules and within the folia at different locations and depths. Total PC numbers and percentage of abnormal PC in different areas within lobules II, III, V (anterior) and lobules VI, VIII, IX (posterior) of the vermis are presented. Total numbers are presented as mean (SD), percentages of abnormal PC are presented as median (IQR).

## DISCUSSION

In this study we investigated PC injury among different locations in the vermis in (near-) term neonates with HIE. We found that both in the folia and the lobules the total PC count was significantly higher at the crowns compared to the bases of the sulci. For several locations the percentage of abnormal PCs was significantly higher at the bases of the sulci than at the crowns. The total number of PCs did not differ between the anterior and posterior lobes of the vermis. To our knowledge this is the first study that describes differences in total PC count within the lobules and folia in the vermis of term born neonates with HIE.

The lower PC count at the bases of the cerebellar lobules and folia compared to the crowns might well be explained by a combination of normal anatomy and increased vulnerability (29,30). Regional differences in PC density within the cerebellar folia have been shown previously in healthy human adults and 75-day-old rats (29,30). Both studies found more PCs at the crowns compared to the bases. These studies showed that the apparent differences in PC density were caused by the folding of the cortex; this could also (partly) be an explanation for the difference in total PC number in our population (29,30).

In general, our data showed more abnormal PC soma morphology at the bases compared to the crowns. Hence it seems that PCs at the bases of the folia and lobules are more vulnerable to hypoxia. Likewise, Akima *et al*. showed that PCs at the bases of the sulci were more prone to severe ischemic injury in humans between 0 and 89 years of age (31–33). They hypothesized that the vascular architecture of the cerebellar cortex provided by the meningeal arteries could be an explanation for selective PC vulnerability to hypoxia (31,32). Although this hypothesis remains to be tested, they did describe that the bases of the deeper secondary and tertiary sulci are perfused by very small branches from the larger arteries (33). Because the arterial ramifications at the bases of the sulci are smaller than the arteries at the crown, this might partially explain the higher prevalence of infarctions in the bases of the sulci after severe generalised ischemia (33).

The current findings seem remarkably similar to a particular pattern of supratentorial injury called ulegyria, first described by Bresler in 1899 (34). Bresler identified narrowing of the supratentorial cortical gyri because of scar formation in the brain of a mentally impaired patient, which he called ulegyria (34). Ulegyria is defined by atrophy of the deeper sulcal portions of the cerebral cortex, thereby sparing the crown, and is most pronounced in the watershed regions of the three major cerebral vessels (35–38). More recently, ulegyria has also been described in three infants with intrauterine asphyxia scanned at seven to eight days after birth by Volpe and Pasternak (38). Ulegyria is a hallmark of perinatal asphyxia in term born neonates with HIE, yet it is not exclusive for infancy (36,37). The acute phase of ulegyria is characterized by extensive neuronal loss and signs of hypoxic-ischemic injury at postmortem histopathological stainings, such as neuronal vacuolization, chromatolysis and an influx of macrophages in the bases of the sulci (39). Thereafter, ulegyria is characterized by atrophic changes and gliosis of the subcortical white matter, eventually leading to the so called ‘mushroom gyri’ with a preserved crown on top of a thin stalk of fibrous glial scar tissue (35,37,39). Ulegyria might be caused by distinctions in cortical perfusion, the crown of the gyri is better perfused than tissue at the bottom of the sulci, thereby causing the typical ‘mushroom shaped gyri’ after hypoxic events (40). Even though, to our knowledge, ulegyria has not yet been described in the cerebellum, the pattern of extensive PC loss at the bases of the sulci in the vermis (vermian ‘ulefolia’) seems relatively comparable to ulegyria.

In the present study, no significant differences were found in total PC count between the anterior and posterior lobes of the vermis. Nevertheless, previous studies have shown that the extent of PC death is not uniform over the cerebellum. PC susceptibility to ischemia is partly dependent on whether PCs express zebrin II or not. Zebrin negative PCs were more vulnerable for hypoxic-ischemia in rodent models (26). Furthermore, there is variability in vulnerability of PCs between lobules; PCs in lobules IX and X seem more resistant to death (26). Additional research is needed to investigate differences in the vulnerability of different lobules in the vermis of neonates with HIE.

There were some limitations to this study. The first limitation was the small research population (n=22). This diminished the power of the study and therefore we were not able to investigate the relationship between timing of death and PC injury. The second limitation was that terminal events after withdrawal of care, such as hypoxic events, could have led to additional injury to H&E stained sections of the vermis (18). Due to the variability between death and tissue collection the phenotype scoring system might have been influenced by post-mortem timing. Furthermore, it was not feasible to determine what the normal distribution of PCs is in term born infants due to the absence of a control group and only little variability in the severity of hypoxic-ischemia in this cohort. Lastly, several neonates were treated with therapeutic hypothermia before withdrawal of care to prevent further brain injury. In this study, hypothermia-treated neonates were not separately investigated from non-hypothermia treated neonates, since the latter was just a minor group (n = 5). The severity of cerebellar injury could potentially differ between infants who were treated with hypothermia and those who were not.

Additional research is essential to better understand the causes and consequences of ulegyria in the vermis. It must be determined whether ulegyria injury is restricted to the vermis or if this phenomenon is also present in the cerebellar hemispheres. Also, the influence of the pattern of PC injury on neurocognitive outcome should be investigated. Supratentorial ulegyria seems to be associated with epilepsy (41), but the effect of ulefolia on outcome is unknown. In addition, it would be interesting to further quantify pathophysiology of immune responsive cells, oligodendrocytes and all neuronal cell types and establish a potential link to cerebellar ulegyria. Another potential avenue of clinical research would be to investigate subcortical white matter injury in the cerebellum, since subcortical white matter atrophy is an essential hallmark of supratentorial ulegyria.

## CONCLUSION

In conclusion, we demonstrated that PCs of term neonates with HIE are injured. HIE infants have fewer PCs at the bases of the cerebellar folia and lobules compared to the crown. In addition, PCs at the bases seem to be more vulnerable to hypoxia. This pattern of PC injury is comparable to supratentorial ulegyria.

